# Negative Binomial Mixture Model for Identification of Noise in Antigen-Specificity Predictions by LIBRA-seq

**DOI:** 10.1101/2023.10.13.562258

**Authors:** Perry T. Wasdin, Alexandra A. Abu-Shmais, Michael W. Irvin, Matthew J. Vukovich, Ivelin S. Georgiev

## Abstract

**Motivation:** LIBRA-seq (linking B cell receptor to antigen specificity by sequencing) provides a powerful tool for interrogating the antigen-specific B cell compartment and identifying antibodies against antigen targets of interest. Identification of noise in LIBRA-seq antigen count data is critical for improving antigen binding predictions for downstream applications including antibody discovery and machine learning technologies.

**Results:** In this study, we present a method for denoising LIBRA-seq data by clustering antigen counts into signal and noise components with a negative binomial mixture model. This approach leverages the VRC01 negative control cells included in a recent LIBRA-seq study(Abu-Shmais *et al*.) to provide a data-driven means for identification of technical noise. We apply this method to a dataset of nine donors representing separate LIBRA-seq experiments and show that our approach provides improved predictions for in vitro antibody-antigen binding when compared to the standard scoring method used in LIBRA-seq, despite variance in data size and noise structure across samples. This development will improve the ability of LIBRA-seq to identify antigen-specific B cells and contribute to providing more reliable datasets for future machine learning based approaches to predicting antibody-antigen binding as the corpus of LIBRA-seq data continues to grow.

**Availability and Implementation:** Jupyter notebooks detailing model fitting and figure generation in Python are available at https://github.com/perrywasdin/mixture_model_denoising.

**Supplementary Information:** Supplementary figures are provided in the attached PDF.

## 1. Introduction

B cells are a critical component of the adaptive immune system. They bind to foreign antigens via the B cell Receptor (BCR) in order to form immunological memory or differentiate into antibody-secreting plasma cells.(Inoue and Kurosaki 2023) The B cell compartment within humans is highly diverse, resulting in a repertoire of BCRs capable of exquisite specificity for an almost unlimited number of antigens. The identification of antigen-specific B cells is of interest for the development of vaccines and monoclonal antibody therapeutics, and also provides a basis for improving our understanding of the role of BCR diversity in the adaptive immune response to infection and vaccination. LIBRA-seq (linking B cell receptor to antigen specificity by sequencing) provides a powerful tool for interrogating the antigen-specific B cell compartment and identifying antibodies against antigen targets of interest.(Setliff *et al*. 2019) In this technology (Fig. 1), a panel of antigens are labeled with oligonucleotide barcodes and then mixed with donor peripheral blood mononuclear cells (PBMCs). Antigen-bound B cells are then isolated and sequenced, enabling high-throughput mapping of single B cells to their cognate antigens.

**Figure 1.**
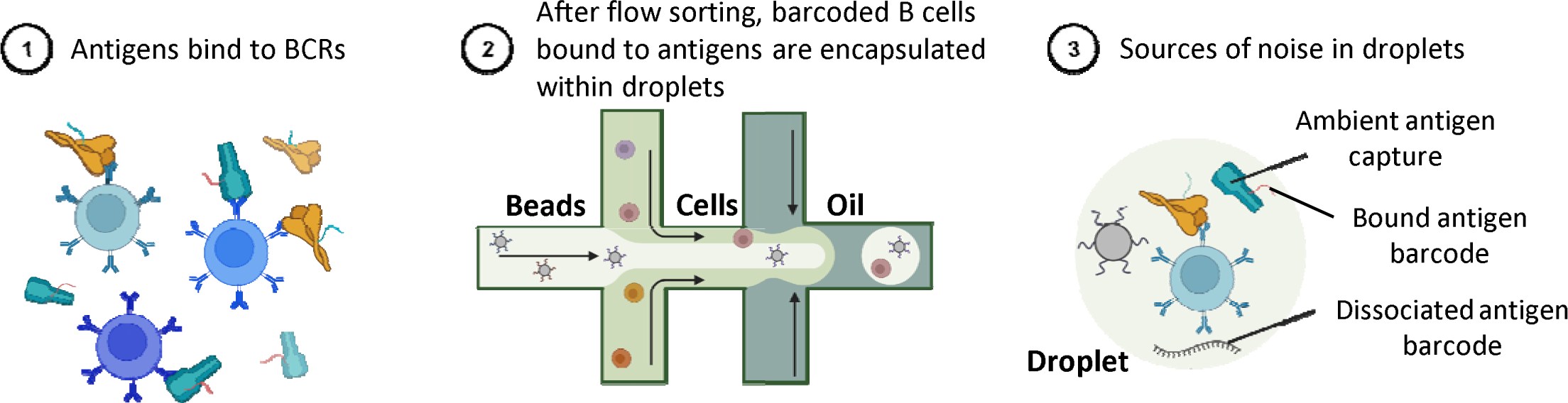
Schematic of LIBRA-seq and sources of technical noise. a) Basic schematic of single cell capture in LIBRA-seq and potential sources of noise in UMI values for an antigen. 1) Barcoded antigens are mixed with donor PBMCs, then antigen-positive B cells are sorted out for sequencing. 2) Droplet encapsulation of single cells and bound antigens. 3) Ambient barcode and antigen capture within droplet lead to false positive UMI counts.

In single-cell sequencing (using the 10X Genomics platform), each molecule in a droplet following the encapsulation of cells (step 2 in Fig. 1) is labelled with a unique oligonucleotide molecular identifier (UMI) tag.(Zheng *et al*. 2017). In the following discussion, UMI counts refer to the UMIs associated with the barcodes used to tag each antigen type in the LIBRA-seq study. The standard LIBRA-seq bioinformatic pipeline uses these UMIs to calculate LIBRA-seq scores (LSS), which represent binding between an antigen and a BCR, based on the number of antigen barcodes associated with each droplet. LSS are calculated by applying a centered log-ratio transform followed by a Z-score transformation to provide relative scores which can be compared across cells for each antigen within a sample. This transformation has been shown to be effective for identification of antigen-specific B cells, but LSS can fail to account for noise due to inflated UMI counts in some LIBRA-seq datasets, resulting in false positive errors for binding predictions. It will therefore be of value to develop an approach for deconvoluting noise from signal in the LIBRA-seq results that can lead to reduced levels of false positive errors.

The denoising approach described here aims to address two major sources of experimental noise which can lead to the superficial inflation of antigen UMI counts (Figure 1, step 3). First, barcodes could dissociate between the final purification step during sample processing and sequencing, resulting in ambient barcodes which are encapsulated in droplets without their antigen. Second, antigens could bind to a B cell or BCR by non-specific binding due to hydrophobic patches, here referred to as “sticky” antigens. While these UMI counts may reflect interaction between a B cell and antigen, they likely do not reflect the highly specific binding interactions which are of interest for therapeutic applications. It is also possible that free-floating antigens in solution could be ambiently captured without interacting with the BCR at all. We do not attempt to differentiate between these processes here, rather we are attempting to identify UMI counts resulting any sources of technical noise that may be present.

Despite these limitations, LIBRA-seq has proven to be immensely useful for antibody discovery applications(Kramer *et al*. 2022; Shiakolas *et al*. 2022; Walker *et al*. 2022; Pilewski *et al*. 2023), and is also quickly becoming a valuable source of antibody-antigen specificity data for computational applications, which has, until recently, been infeasible to collect in a high-throughput manner. While large databases of antibody sequences exist(Abanades *et al*.; Raybould *et al*. 2021; Olsen, Boyles and Deane 2022), there are currently no antigen-specific antibody databases of adequate size and scope for machine learning applications, and synthetic data often must be incorporated.(Akbar *et al*. 2022) The available body of LIBRA-seq data continues to grow, promising to become an important source of data for future technologies aimed at predicting antigen specificity. However, the usefulness of this data for such applications hinges on the reliability of binding predictions. Aside from machine learning applications, further experimental validation of antigen binding predictions from LIBRA-seq is expensive and time consuming, and the field stands to benefit greatly from improved accuracy of LIBRA-seq antigen specificity predictions.

## 2. Methods

### 2.1 Datasets

A recent study from our group aimed to generate large-scale antibody-antigen datasets, detailing the collection and characterization of LIBRA-seq data for a set of 10 healthy donors (samples) across a panel of 20 antigens.(Abu-Shmais *et al*.) This study provides a rich dataset for development of the denoising model presented here. In the LIBRA-seq experiments, an antigen screening library of 20 antigens from diverse but common pathogens were included. In addition, cells from the Ramos cell line which were engineered to express the HIV-1-specific antibody VRC01(Zhou *et al*. 2010) were also included, along with an HIV-1 antigen (BG505), to serve as negative controls: since the donors were screened for HIV-1, they theoretically should have very few antibodies which bind to the BG505 antigen, while the VRC01 expressing cells should only bind to the BG505 antigen and none of the other antigens in the LIBRA-seq screening library. In our approach, we leveraged these negative control HIV-1 specific cells by assuming that UMI counts associated with VRC01 cells and non-HIV antigens represent technical noise in the experiment. As a result, the VRC01 cells provide information about the structure of noise for each antigen in each sample and could be used to bias our denoising algorithm. In vitro validation data was also collected for 99 antibodies of interest, using their BCR sequences to recombinantly express the antibodies and test for binding against their identified cognate antigen using ELISA (enzyme-linked immunosorbent assays).

#### Preprocessing

In the Atlas experiments, PBMCs from each donor were processed and sequenced as individual samples. Raw sequencing reads from 10X single cell RNA- and VDJ-sequencing were aligned to the human reference genome GRCh20 (2020) using Cell Ranger 3.1.0. Calculation of LIBRA-seq scores (LSS) was performed using the LIBRA-seq bioinformatic pipeline, as previously described. In the standard pipeline, UMI counts <4 are typically set to 0 before the LSS transformation, as UMI counts this low are considered unreliable. For the denoising approach presented here, these UMI counts were retained so that the count distributions would accurately reflect the structure of noise and signal captured in the experiments. Cells associated with more than one heavy chain (N>1) were removed before denoising as these cells could potentially represent sequencing errors or multiplet capture. For each sample, VRC01 cells were separated from the real donor cells by comparing the heavy chain CDR3 sequences to the VRC01 sequence with a Levenshtein distance threshold of 0.05 (representing 95% identity). UMI counts > 99^th^ percentile were assumed to be outliers and were removed from the VRC01 and donor B cell distributions.

Cell counts following this preprocessing are summarized in Table 1 for nine donors chosen to be used in the development and application of our denoising approach here. The total number of donor B cells recovered ranged from 160 B cells in Donor 5 to 4,100 B cells in Donor 1. The number of negative control cells recovered ranged from 0 cells in Donor 3 to 6,766 cells in Donor 6. Donor 2 was chosen to be used for example visualizations in the Results section as a substantial number of both donor and VRC01 cells were recovered, but other donors are included in the figures and supplement as applicable. Despite the broad antigen panel used in the experiments, SARS-CoV-2 spike was the only antigen consistently recovered in high numbers across donors. An average of 486 UMI counts were recovered for SARS-CoV-2 across all donors, compared to an average of less than 50 cells with >10 UMI counts for all other antigens. Additionally, some donors from the original set of experiments were excluded based on low cell counts (<50 total) and overall poor sample quality. Further insight into the necessary data size for this approach is included later in the discussion.

**Table 1.** Summary of cell counts across all donors. Cell counts shown for donor cells with >10 SARS-2 UMI counts, total donor B cells, and total VRC01 Ramos negative control cells following preprocessing.

| **Cells** | **Donor ID** |  |  |  |  |  |  |  |  |
| --- | --- | --- | --- | --- | --- | --- | --- | --- | --- |
|  | **1** | **2** | **3** | **4** | **5** | **6** | **7** | **8** | **9** |
| **SARS-CoV-2** | 253 | 1182 | 4712 | 48 | 39 | 276 | 356 | 215 | 181 |
| **Donor Total** | 4100 | 1690 | 4721 | 229 | 160 | 523 | 361 | 226 | 199 |
| **VRC01** | 43 | 1155 | 0 | 357 | 351 | 6766 | 3424 | 139 | 214 |

#### Characterization of noise in data

Comparing the distribution of antigen UMI counts in donor B cells to VRC01 negative control cells provides a clear visualization of noise for a given sample. In the donor cells (Fig. 2a), a bimodal distribution is seen with one component of lower counts and a second distribution with larger variance at higher UMI counts. In VRC01 cells from the same sample (Fig. 2b), there is a single distribution which appears similar to the lower count distribution observed in the donor cells, suggesting that the antigen UMI counts in the VRC01 cells are representative of noise in the donor cells. The second component in the Donor cells, which is not recapitulated in the negative control, is assumed to represent true signal from binding interactions between the B cells and antigen.

**Figure 2.**
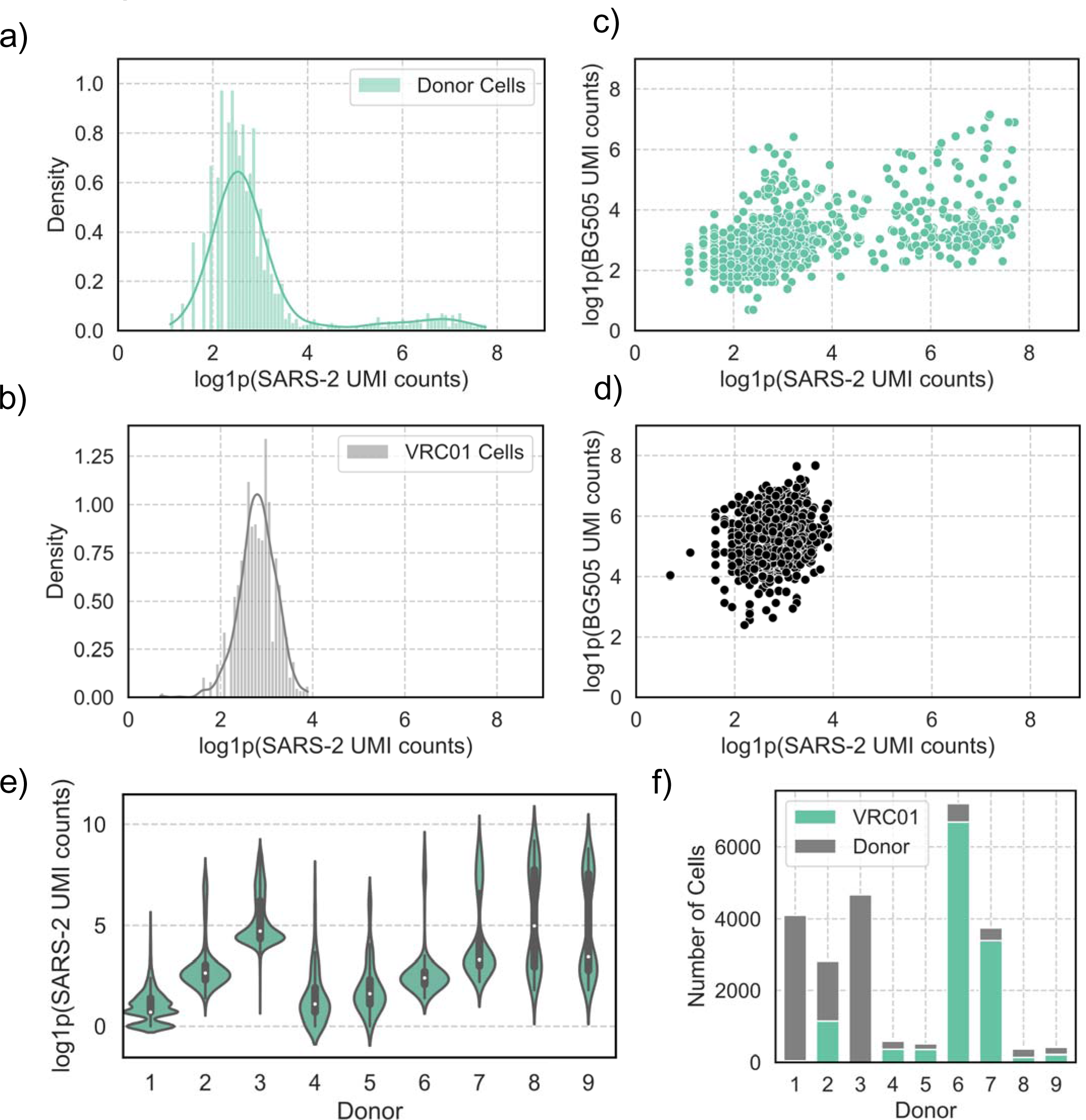
Characterization of UMI noise LIBRA-seq data. Histograms with kernel density estimates (KDEs) representing distributions of SARS-CoV-2 spike (SARS-2) UMI counts for a) donor B cells and b) VRC01 negative control cells in Donor 2. c) Plotting BG505 (HIV-1) UMI counts against SARS-CoV-2 UMI counts for Donor 2. d) BG505 (HIV-1) UMI counts against SARS-CoV-2 UMI counts for Donor 2 VRC01 cells. e) Violin plot comparing distributions of SARS-CoV-2 UMI counts for B cells across donors. f) Stacked counts of donor B cells and VRC01 cells for each donor.

A similar bimodal distribution is also seen in the relationship between the SARS-CoV-2 UMI counts and the UMI counts for BG505, which serves as a negative control (Fig. 2c). For Donor 2, there is a positive correlation (Spearman’s r=0.48) between these distributions. In the complete absence of noise, we would expect this correlation to be 0, but this positive correlation implies that there is some degree of non-specific (promiscuous) binding between the BCRs and antigens as very few BCRs should bind to both HIV-1 Env and SARS-CoV-2 spike. The VRC01-expressing negative control cells exhibit more specificity, with a Spearman correlation of 0.21 between BG505 and SARS-CoV-2, suggesting that the BCRs may be responsible for non-specific binding rather than antigen stickiness. While we do not attempt to explore the source of this noise or the binding to BG505 any further in this manuscript, the relationship between SARS-CoV-2 and BG505 UMI counts further motivates the need to filter out false positive UMI counts. Despite the lower correlation between these UMI count distributions for VRC01 cells, Figure 2d emphasizes that even with an antibody well known to be highly specific to HIV-1, there is a distribution of inflated UMI counts captured for SARS-CoV-2 that represents technical noise in the sample.

The structure of UMI signal and noise is complicated by the variance of distributions across donors (Fig. 2e). For most of the samples, there was a clear visual separation of multiple distributions for SARS-CoV-2 UMI counts (Fig. S1.). The distributions of SARS-CoV-2 UMI counts in VRC01 cells, representing noise (Fig. S2.), also showed variation across donors. This variation implies that a single threshold for UMI signal/noise may not be sufficient for filtering across datasets, emphasizing the need for a more sophisticated approach to identifying noise and establishing biologically meaningful thresholds to indicate binding. Sample-to-sample variation was also apparent in the differences observed in the ratio of donor:VRC01 cells (Fig. 2f). The impact of VRC01 on model fitting and accuracy is explored later, as some donors display very low counts of VRC01, with Donor 3 representing an extreme where zero VRC01 cells were recovered.

### 2.2 Algorithm

The algorithm described here represents a custom mixture model approach using negative binomial distributions to cluster experimental UMI counts. After separating the VRC01 cells from the real donor B cells, the following pipeline was applied to the distribution of counts for each antigen within each donor separately. This approach consists of three steps, which are visualized in Figure 3. First, a maximum likelihood estimator (MLE) was used to fit a negative binomial distribution to the UMI counts associated with the VRC01 negative control cells. The probability mass function for the negative binomial distribution for a UMI count *k* (Eq. 1) is described by the parameters *n* and *p*.(Virtanen *et al*. 2020)

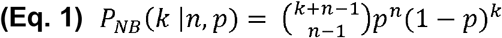

The parameters from the noise distribution are then used to initialize step two, where a negative binomial mixture model is fit to the UMI count distribution for the donor cells (Eq. 2). The mixed distribution consists of two negative binomial components and a weight w to account the imbalanced cell counts between the number of VRC01 cells and B cells for different donors. The parameters for this mixed distribution are estimated with an MLE, using the log of the sum of exponentials to minimize the negative loglikelihood of the component mixture.

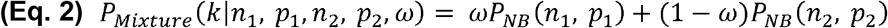

Finally, Bayes’ Theorem was employed using the probability mass functions for each component to calculate the probability for each observation (UMI count) of being in either the signal (P_S_) or noise (P_N_) component (Eq. 3). In the mixed distribution, the components are nonidentifiable, meaning that the component labels are initially arbitrary and are not considered as signal or noise during fitting by MLE. Component labels are assigned post-hoc, where the component with the higher median was considered the ‘Signal’ component and the lower median component was labeled as ‘Noise’. In the results and discussion here, the probabilities referenced will be P_S_, the probability of a given UMI count being in the signal component of the mixed distribution.

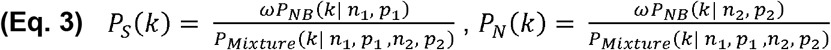

The pipeline was implemented using the *Statsmodels*(Seabold and Perktold 2010) and *Scipy*(Virtanen *et al*. 2020) libraries in Python. For step 1, fitting of the noise distribution P_NB_, the MLE was initialized with started parameters *p = n = 1*. In step 2, fitting of the mixed distribution P_Mixture_, the weight was initialized as w= 0.1. The parameters from the noise distribution were set as *n*_*2*_ and *p*_*2*_, while *n*_*1*_ and *p*_*1*_ were estimated based on the initial weight guess. Briefly, the weight guess was used to split the UMI count data into two components based on counts above and below a percentile calculated from the weight, then a negative binomial distribution was fit to the lower percentile distribution to estimate starting parameters *n*_*1*_ and *p*_*1*_ for initializing the mixed distribution MLE. For Donor 3, where no VRC01 cells were recovered, this percentile-based approach for estimating starting parameters was used to initialize parameters for both components.

**Figure 3.**
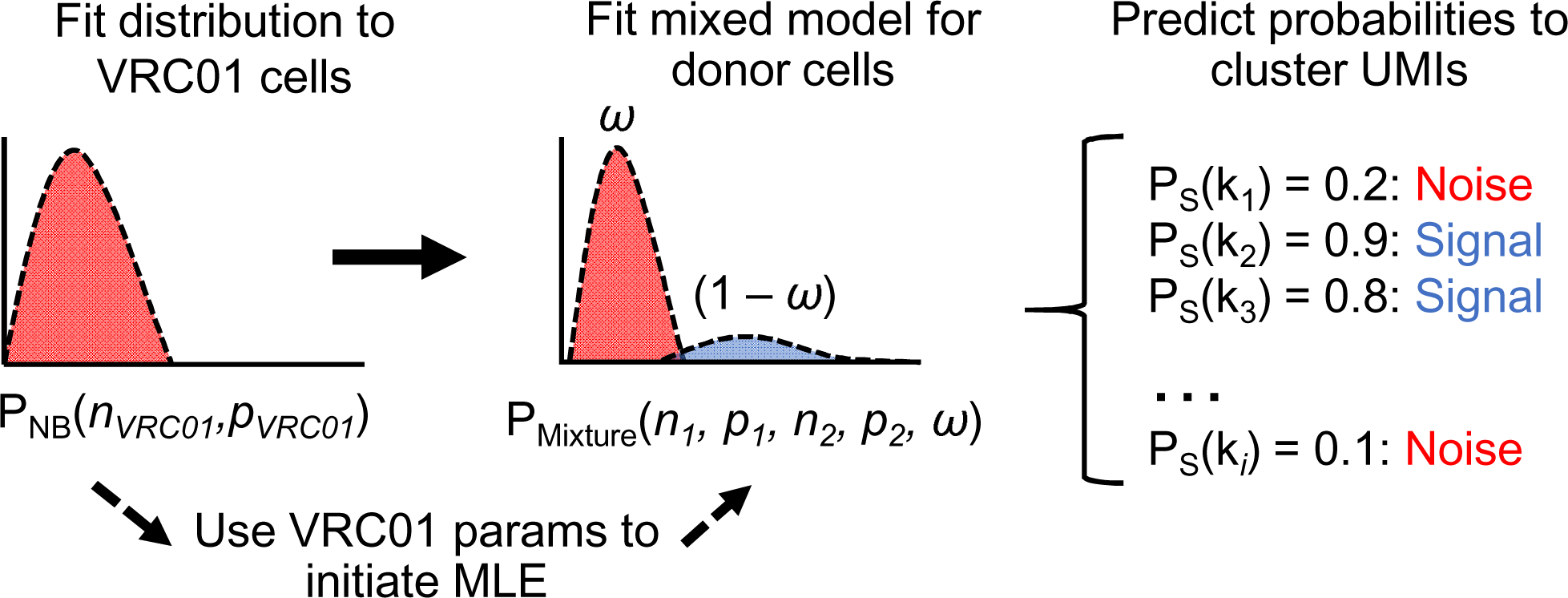
Schematic of mixture model pipeline. First, a distribution is fit to the VRC01 UMI count distribution for SARS-CoV-2 using an MLE. Then, these parameters are used to initialize an MLE to fit a mixed distribution to the SARS-CoV-2 UMI count distribution for the donor B cells. The probability mass function of the mixed distribution is used to generate probabilities for the SARS-CoV-2 UMI counts (k) for all cells (i) in a donor. Finally, the UMI count is then assigned to one of the components based on a threshold for PS and labeled as either ‘signal’ or ‘noise’.

## 3. Results

A critical assumption underlying the use of LSS for binding predictions is that antigen barcodes captured in a droplet represent true binding interactions between the encapsulated B cell and the antigen from which the barcode originated. As discussed earlier, various sources of technical noise can inflate these UMI counts, leading to false positive predictions. The pipeline presented here aims to fit a mixed distribution model to UMI count distributions in the donor cells which can then be used to cluster the counts into noise and signal components, enabling more accurate identification of antigen-specific B cells. This approach is employed separately for each donor and antigen to account for the variability in noise observed across donors.

Counts in single cell data are commonly modeled using a variety of distribution types including Gaussian(Mulè, Martins and Tsang 2022) or discrete distributions such as the Poisson and negative binomial distributions.(He *et al*. 2021; Fleming *et al*. 2023) In order to determine the best approach for modelling noise in the VRC01 cells, we fit three common distribution types (Fig. S3a) and found that the negative binomial distribution (Fig. 4a) had the best fit according to negative loglikelihood (NLL), Akaike Information Criterion (AIC), and Bayesian Information Criterion (BIC). Similarly, after fitting the mixed distributions to the donor cell distributions, we found that the mixed negative binomial model was most representative of the empirical distribution (Fig. 4b) for UMI counts in the donor B cells. The mixed Poisson distribution fit poorly overall, while the mixed Gaussian model appeared to fit very well in the middle range of UMI counts but failed to capture the behavior of the distribution at the extremities for Donor 2. The mixed negative binomial distribution fit with a lower BIC than the mixed Poisson distribution for all donors (Fig. 4d). The mixed Gaussian model was also tested across all donors but consistently failed to converge. (Fig. S3b).

**Figure 4.**
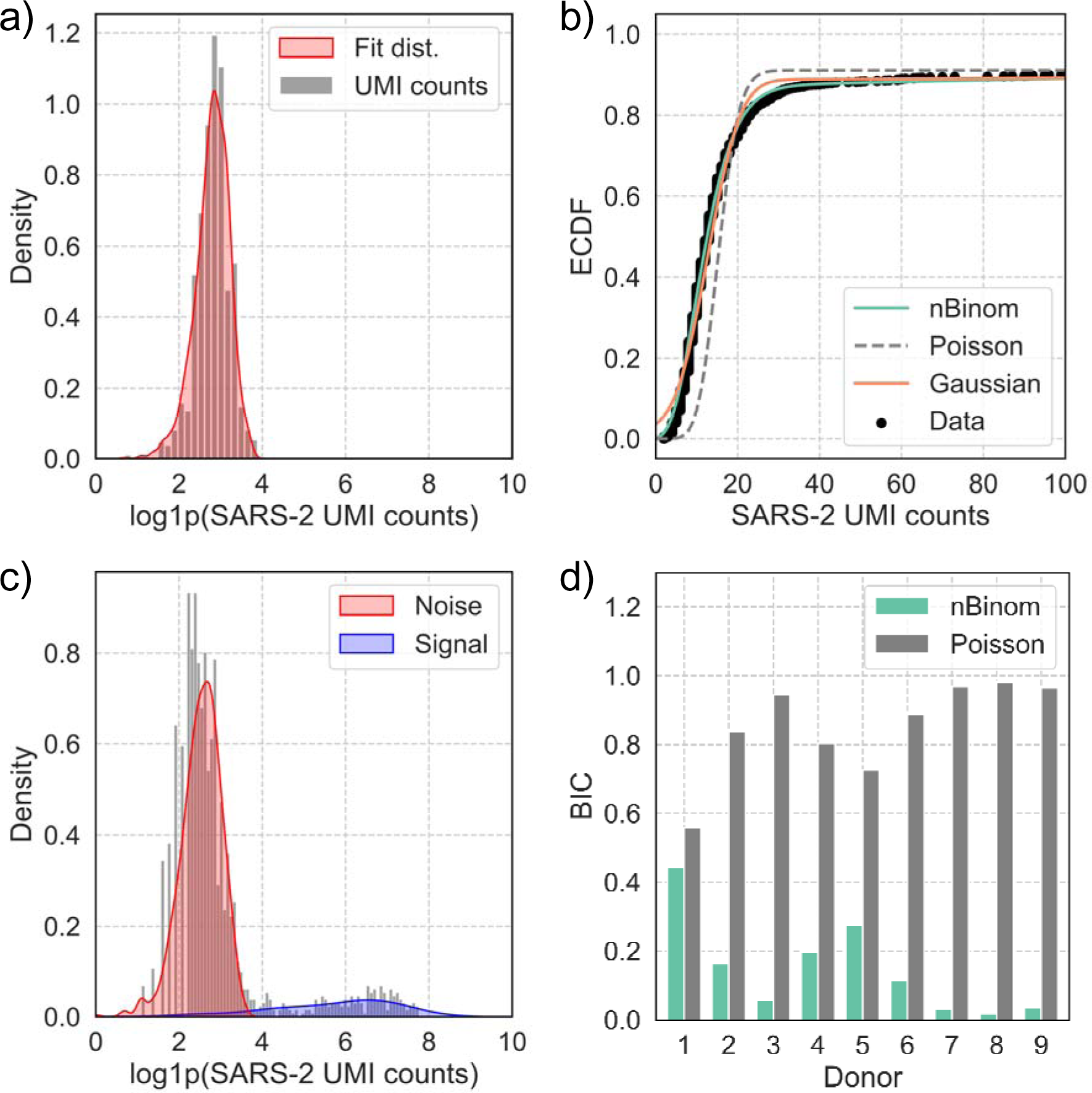
Assessment of fit quality for noise and mixture models. a) Visualization of randomly sampled variates from noise distribution fit to SARS-CoV-2 spike (SARS-2) UMI counts captured for VRC01 cells in Donor 2. b) Comparison of empirical cumulative distribution functions (ECDFs) for mixed distributions of Gaussian, Poisson, and Negative Binomial distributions for Donor 2 SARS-CoV-2 UMI counts. c) Visualization of KDE for the two components fit for the negative binomial mixture model of SARS-CoV-2 UMI counts for Donor 2. d) Comparison of Bayesian Information Criteria (BIC) from mixed model fits for each donor.

In addition to BIC, the fit quality of the mixed negative binomial distributions can also be visually confirmed for the VRC01 noise distribution (Fig. 4a) and the mixture model of the donor B cells (Fig. 4c) for Donor 2, along with the rest of the donors (Fig. S4). These visualizations emphasize some of the challenges in finding a model that can account for the large variations in UMI distributions observed across donors here. Donor 1, for example, appeared to primarily contain noise with low UMI counts and very few true signal counts. Donors 2 and 6-9 showed relatively well separated distributions of low and high UMI counts, corresponding to the noise and signal components.

### Determining quality of signal probability predictions

After fitting the negative binomial mixture model to the donor cell UMI count distribution for SARS-CoV-2 spike, an adaption of Bayes’ theorem was applied to the probability mass function (PMF) of the mixed distribution to calculate the probabilities of each UMI count belonging to the signal component. The probabilities output from Bayes’ theorem can then be binned with a threshold to cluster the UMI counts as ‘signal’ or ‘noise’, similar to a Gaussian mixture model clustering algorithm(Murphy 2012).

Once signal probabilities (P_S_) are obtained, the challenge remains to establish a biologically relevant threshold to predict true binding. While this is straightforward with clearly separated distribution components, determining a clustering threshold is more challenging for donors where the two distributions overlap. An unbiased threshold can be chosen as P_S_ = P_N_ = 0.5, but this threshold could be modified depending on the application. For example, in antibody discovery it may be advantageous to identify a greater number of potential antibody candidates despite the increased risk of false positives, while machine learning applications would benefit from more conservative thresholds to reduce false positives.

In Figure 5a, we can see that a sigmoidal curve is formed when comparing the signal probabilities to the antigen UMI counts. Typically, a LSS threshold of 1 is used to classify binding, based on experimental observations during the initial development of the method. In Figure 5b we see that applying this LSS threshold alone includes a number of probabilities below 0.5 which could result in false positive binding assignments when considering probability alone. Since this is a probabilistic model that aims to identify true UMI counts, rather than directly predicting binding, we recommend using the calculated signal probability in conjunction with the standard LSS threshold to yield the highest confidence predictions. For Donor 2 (Fig. 5c), we can see that the component assignments from clustering intuitively aligned with the anticipated signal/noise division that we would expect based on the VRC01 distribution shown previously in Figure 2.

**Figure 5.**
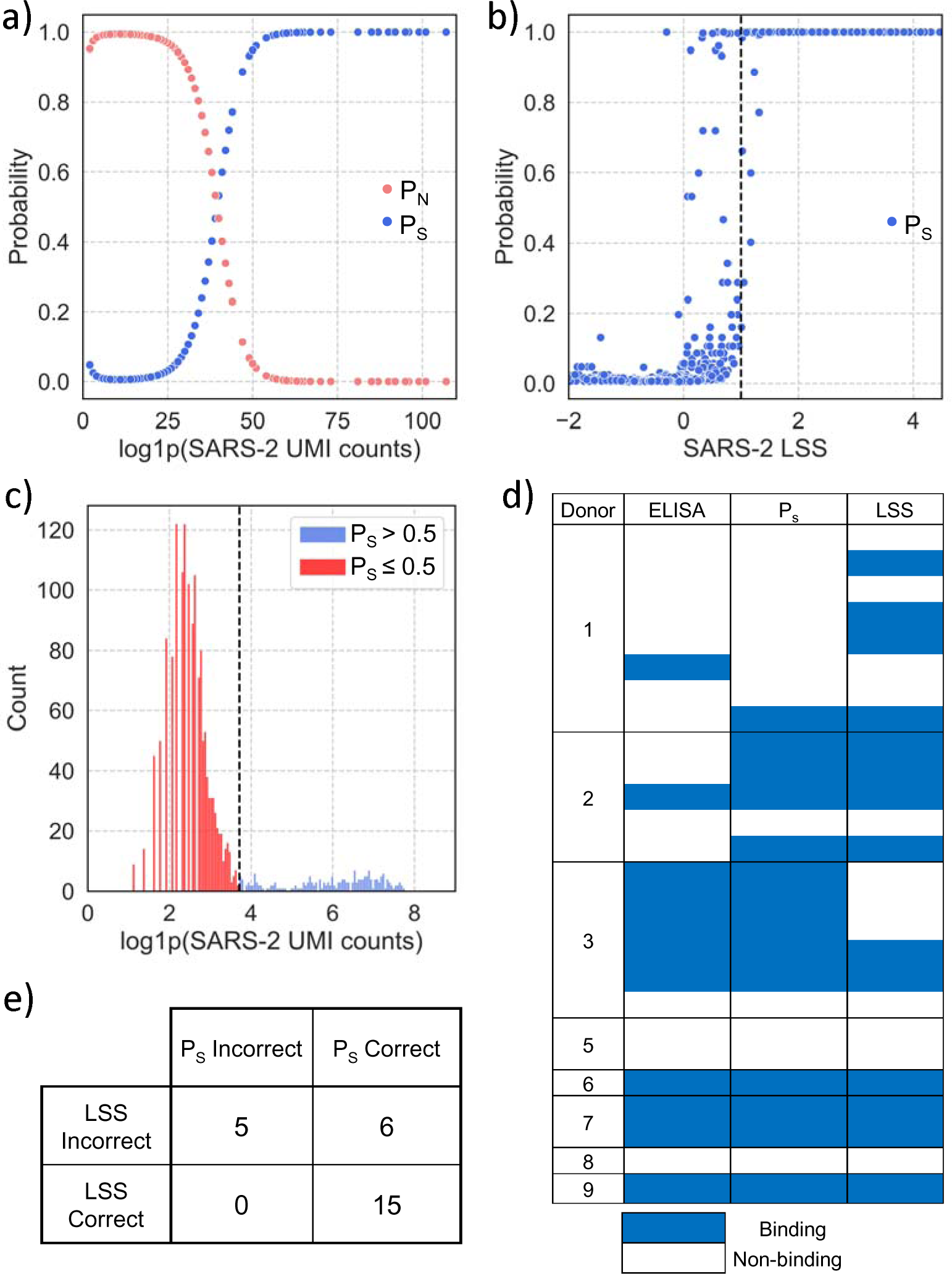
Model predictions and in-vitro validation. After model fitting, the mixture model was used to cluster SARS-CoV-2 spike (SARS-2) UMI counts into the signal and noise components. a) Plotting P_S_ against UMI counts for Donor 2 shows the inflection point between components. b) Plotting P_s_ against LSS for Donor 2 with dotted lines representing the recommended thresholds for each. c) Histogram of log1p transformed UMI counts, with labels assigned based on a P_s_ threshold of 0.9. d) Absorbance (AU_450_) values from ELISA micro-expression for recombinantly expressed antibodies compared to LSS and P_S_ predictions. Binned values are shown based on the following thresholds: LSS ≥ 1, P_S_ ≥ 0.9, AU_450_≥ 1. e) Confusion matrix comparing P_s_ to ELISA values summarizing results from panel d.

### In vitro validation of binding predictions

In the LIBRA-seq study, binding predictions by LSS were tested *in vitro* by recombinantly expressing the antibodies and performing enzyme linked immunosorbent assays (ELISA).(Abu-Shmais *et al*.) We compared these results to predictions made by the signal probabilities (P_S_) generated by the mixture model here to determine the accuracy of the model and see how it compares to the standard LSS. For the donors selected here, there were a total of 26 antibodies validated for binding to SARS-CoV-2 spike. LSS accurately predicted binding for 14/26 (54%) antibodies, while the mixture model accurately predicted binding for 20/26 (77%) antibodies based on P_S_ (Fig. 5d). This increase in performance was mainly due to improved predictions in Donors 1 and 3, while predictions for all other donors matched the LSS. Importantly, there were no instances where the LSS was correct and P_S_ was wrong (Fig. 5e), meaning that the probabilities always matched or improved predictions. In order to predict binding using LIBRA-seq, a fundamental assumption is that UMI counts represent binding, although this relationship may not always be linear or correlative. In many cases where both LSS and P_S_ incorrectly predicted, the UMI counts do not appear linked to binding at all, with one example where an antibody bound in ELISA despite only yielding 3 UMI counts for SARS-CoV-2 spike. Correcting discrepancies such as this is outside the scope of this model, as these are clearly not due to UMI count inflation alone and could result from technical issues during any of the experimental steps in the pipeline.

## 4. Discussion

Identification of noise in LIBRA-seq UMI count distributions is critical for improving antigen binding predictions for downstream applications such as antibody discovery and machine learning technologies. In this study, we present a method for clustering antigen UMI counts into signal and noise components with a negative binomial mixture model. This approach provides improved antigen binding predictions across multiple samples when combined with the standard scoring method (LSS), despite heterogeneity in data size and noise structure.

The pipeline described here leverages the VRC01 negative control cells included in the LIBRA-seq study(Abu-Shmais *et al*.), providing a data-driven means for introducing bias to model fitting. While this negative control was important for understanding the structure of noise in this dataset, the inclusion of such cells increases cost and labor and can also detract from the total recovery rate of donor B cells for each sequencing run. The Ramos cell line can also be sensitive to handling and recovery can be poor, as seen here with Donor 1 only containing 43 VRC01 cells, and Donor 3 recovering zero. Additionally, such controls were not included in previously published LIBRA-seq experiments and could limit the application of this method. However, we have shown that the mixture model for noise clustering was still able to enhance LSS predictions for Donor 3, where no VRC01 cells were recovered. To further support the possibility of using this approach in the absence of VRC01, the pipeline was tested on all donors without inclusion of bias from VRC01 for the mixture model MLE. For this approach, the mixed distribution parameters were initialized using the strategy previously described for Donor 3, where mean and variance were estimated from an initial weight guess. Here, P_S_ yielded the same predictions as before for Donors 1-5 but performed worse on Donors 6-9. This suggests that the mixture model approach may work well in the absence of VRC01 with adequate data size but can lose accuracy for donors with limited available data.

Another limitation of the mixture model denoising approach described here is the data size required for accurate clustering. The size threshold for choosing samples initially was somewhat arbitrary since it is difficult to quantify the exact number of cells needed without a much larger in vitro validation dataset. In an attempt to estimate the necessary data size for our approach to work, bootstrapping with downsampling was performed (Fig. S5). Donor cells were downsampled across a range of total B cell counts in order to estimate model stability. As shown in Figure S5a, the coefficient of variation for the BIC of the mixed distribution fit begins to increase rapidly for most donors once the sample size drops below 300 total donor B cells, suggesting a decrease in model fit at low sample sizes. Similarly, the variation of P_S_ increases below 300 cells, showing an increased uncertainty in model predictions (Fig. S5b). Based on this bootstrapping analysis, Donor 1 appeared to be most sensitive to decreased sample size. While this may be surprising since Donor 1 had high overall cell counts, it also had the lowest proportion of cells with SARS-CoV-2 UMI counts > 10 (7.14%). Due to the previously discussed variation is UMI count distributions, the necessary sample size may ultimately depend on the separation and ratio of noise/signal components, but bootstrapping suggests that a total cell count of at least 300 B cells is necessary for stable model fitting and low prediction error.

The standard error of the mean signal probabilities for 100 iterations of bootstrapping with resampling are shown in Figure S5c. Donor 1 displayed the highest error of 2.2% SEM, likely due to the low ratio of high SARS-CoV-2 UMI counts relative to the noise distribution, as mentioned above. For all other donors, the error in P_S_ was <1%, suggesting that the model fit reliably and was not sensitive to cell sampling. Overall, we have shown that our negative binomial mixture model approach for denoising LIBRA-seq data is able to accommodate sample-to-sample variability and improve antigen-binding predictions for B cells across 9 individual samples collected from separate donors. This development will improve the ability of LIBRA-seq to identify antigen-specific B cells and contribute to providing more reliable datasets for future machine learning based approaches to predicting antibody-antigen binding as the corpus of LIBRA-seq data continues to grow.

## Supporting information

All supplemental figures

## Acknowledgements

We thank all members of the Georgiev lab for their support and feedback. This work was conducted, in part, using the resources of the Advanced Computing Center for Research and Education (ACCRE) at Vanderbilt University. For the work described in this manuscript, I.S.G., P.T.W, A.A.A., and M.J.V. were supported in part by the G. Harold and Leila Y. Mathers Charitable Foundation (MF-2107-01851), NIH R01AI175245 (to I.S.G.), and NIH R01AI152693-03 (to I.S.G.). A.A.A. was supported in part by NIH grant T32 (5T32AI112541-07) and P.T.W. was supported in part by NIEHS grant T15 (T15LM007450-19). The funders had no role in the conceptualization or execution of any studies or drafting of the manuscript.

## Author Contributions

Conceptualization and Methodology: P.T.W., M.W.I., and I.S.G.; Investigation: P.T.W, A.A.A., M.J.V.; Writing – Original Draft: P.T.W. and I.S.G.; Writing – Review & Editing: All authors; Funding Acquisition: P.T.W, A.A.A., and I.S.G. Resources: I.S.G; Supervision: P.T.W and I.S.G

## Declaration of Interest

I.S.G. is listed as an inventor on patent applications for the LIBRA-seq technology. I.S.G. is a co-founder of AbSeek Bio. I.S.G. has served as a consultant for Sanofi. The Georgiev laboratory at VUMC has received unrelated funding from Merck and Takeda Pharmaceuticals.

## Figure Titles and Legends

**Figure S1. SARS-CoV-2 UMI count distributions across donors**

Histograms representing distributions SARS-CoV-2 spike (SARS-2) UMI counts captured for donor B cells in each donor.

**Figure S2. SARS-CoV-2 UMI count distributions for VRC01 across donors**

Histograms representing distributions SARS-CoV-2 spike (SARS-2) UMI counts captured for VRC01 expression Ramos cells in each donor.

**Figure S3. Fit quality metrics for SARS-CoV-2 UMI distributions across donors**

a) Comparison of BIC for negative binomial, Poisson, and Gaussian distributions fit to SARS-CoV-2 spike (SARS-2) UMI counts in VRC01 cells. b) Comparison of BIC for mixture models of binomial, Poisson, and Gaussian distributions fit to SARS-CoV-2 UMI counts in donor cells.

**Figure S4. Mixed model fit for all samples for SARS-2**

a) Visualization of KDE for the two components fit for the negative binomial mixed distributions of SARS-CoV-2 spike (SARS-2) UMI counts for all 9 donors.

**Figure S5: Bootstrapping and uncertainty quantification**

a) Line plot representing coefficient of variance for BIC of mixed distribution fit across series of downsampled datasets for Donor 2 SARS-CoV-2 UMI counts in each donor. 50 iterations were performed per downsampled data set per donor, with resampling. b) Coefficient of variance for standard error of the mean signal probability (P_S_) across all cells at each sampling iteration. c) Mean percent standard error across all cells for each donor after 100 iterations of bootstrapping with resampling.

**Figure S6. Model can work in absence of negative control with adequate amount of data**

Comparison of ELISA Absorbance (AU_450_) values from ELISA micro-expression for recombinantly expressed antibodies with P_s_ and LSS predicting binding to SARS-CoV-2 spike using the mixture model fit to all donor cells for each donor, without inclusion of VRC01. Binned values are shown based on the following thresholds: LSS ≥ 1, P_S_ ≥ 0.9, AU_450_≥ 1.

## Notes

### Summary of Updates

Revised labels on Figure 5; added Figure numbers.

