## Supplementary figures and images for "Negative Binomial Mixture Model for Identification of Noise in Antigen-Specificity Predictions by LIBRA-seq"

### All supplemental figures

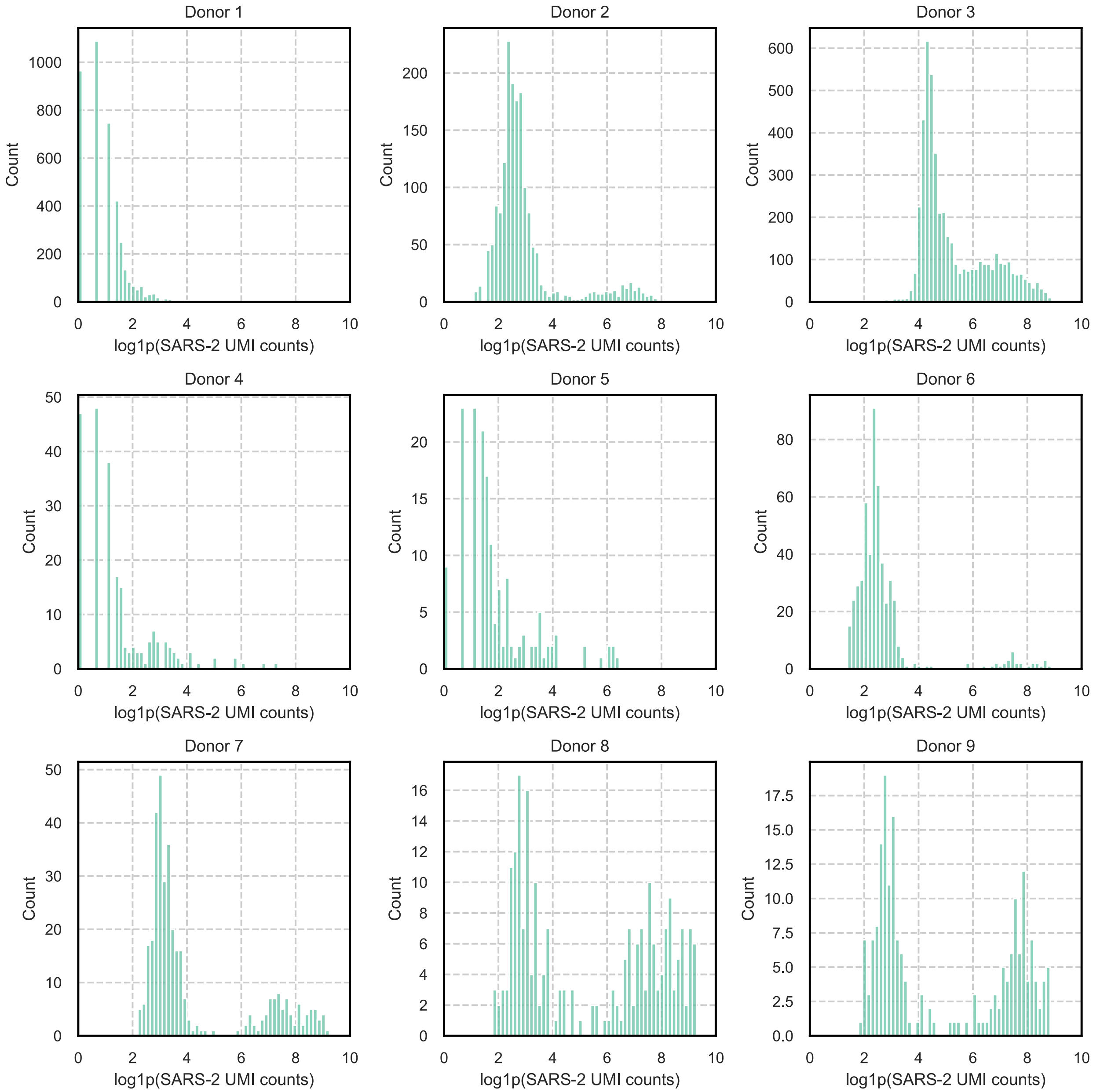

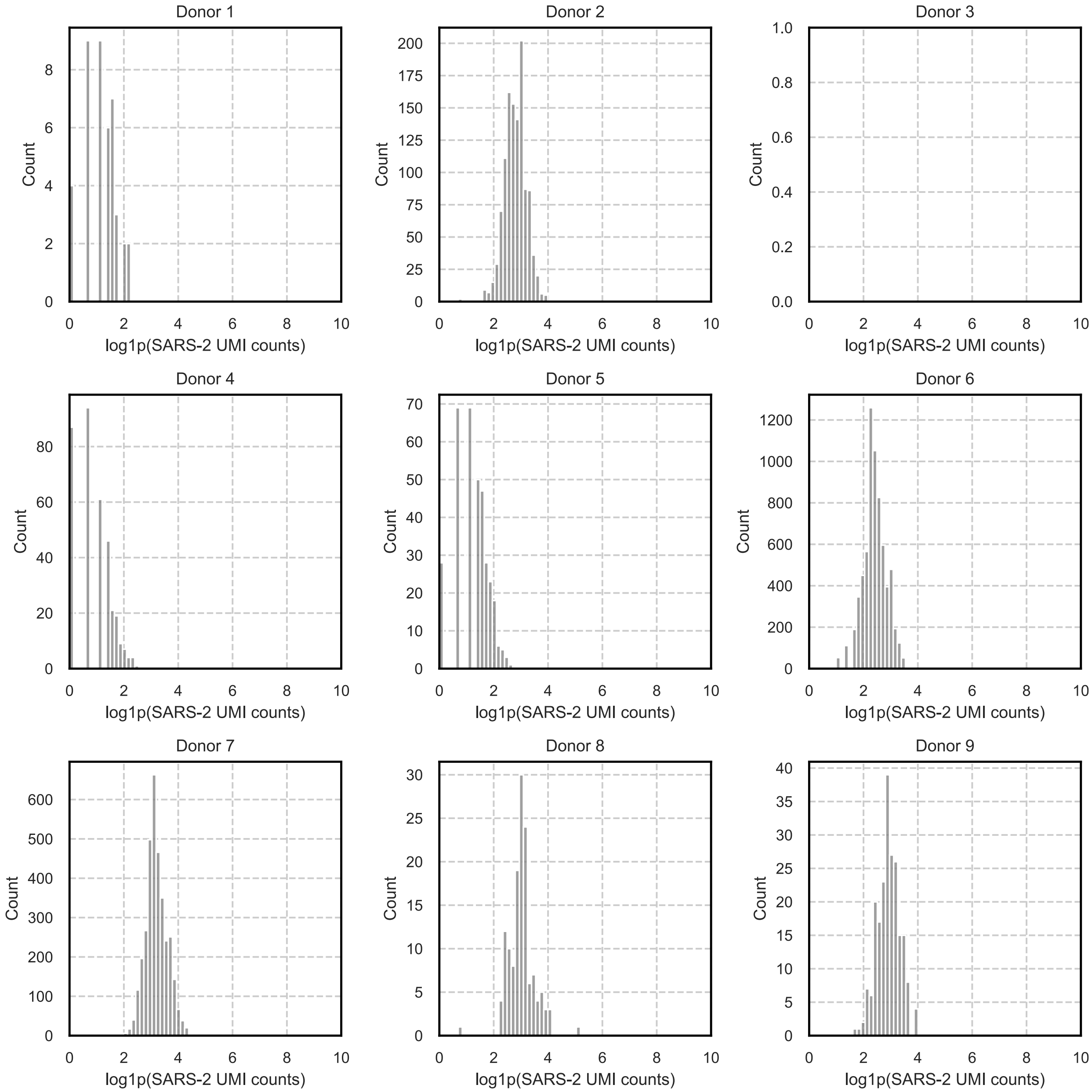

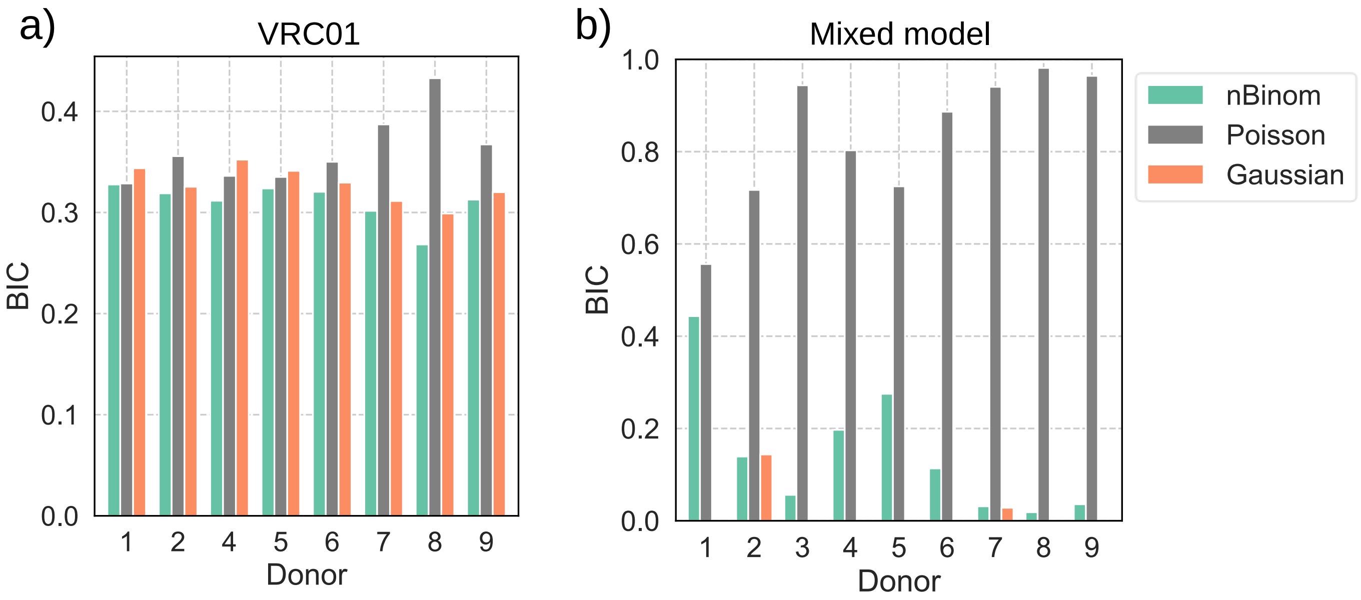

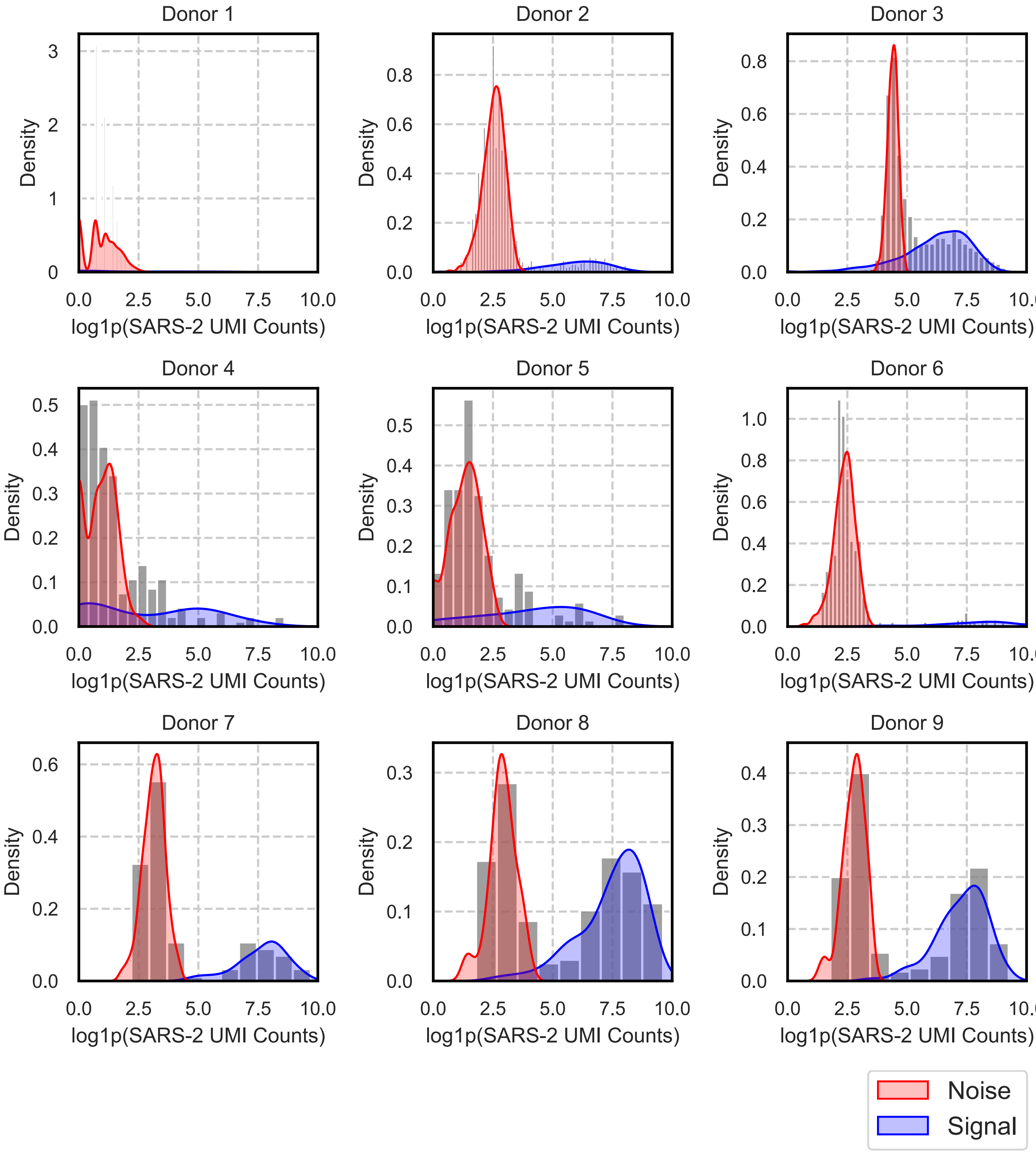

a)

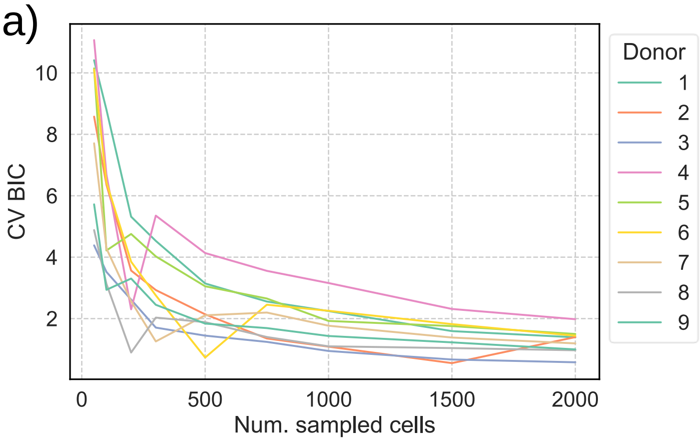


b)

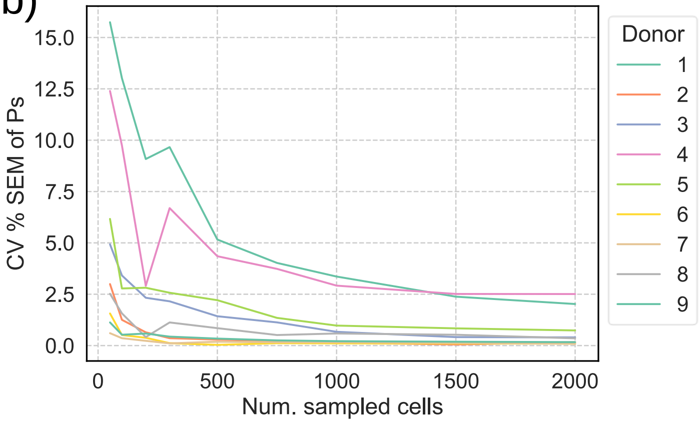


c)

| Donor | Ps SEM (%) |
|---|---|
| 1 | 2.239 |
| 2 | 0.022 |
| 3 | 0.022 |
| 4 | 0.406 |
| 5 | 0.232 |
| 6 | 0.048 |
| 7 | 0.081 |
| 8 | 0.071 |
| 9 | 0.071 |

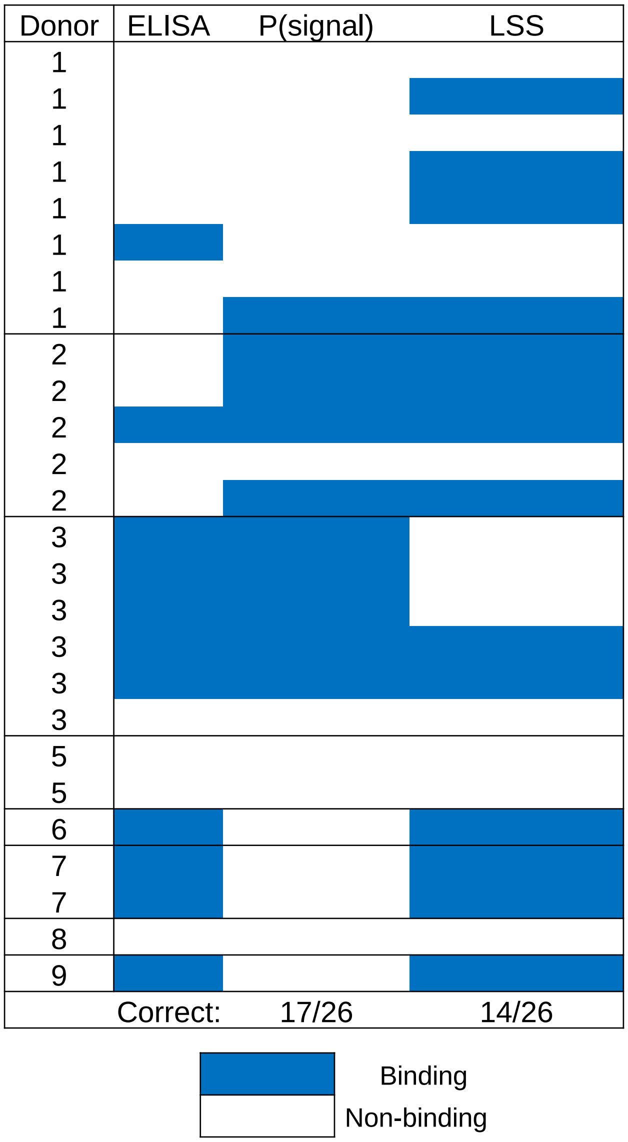
